# A network of epigenomic and transcriptional cooperation encompassing an epigenomic master regulator in cancer

**DOI:** 10.1101/309484

**Authors:** Stephen Wilson, Fabian V. Filipp

## Abstract

Coordinated experiments focused on transcriptional responses and chromatin states are well-equipped to capture different epigenomic and transcriptomic levels governing the circuitry of a regulatory network. We propose a workflow for the genome-wide identification of epigenomic and transcriptional cooperation to elucidate transcriptional networks in cancer. Gene promoter annotation in combination with network analysis and sequence-resolution of enriched transcriptional motifs in epigenomic data reveals transcription factor families that act synergistically with epigenomic master regulators. A close teamwork of the transcriptional and epigenomic machinery was discovered. The network is tightly connected and includes the histone lysine demethylase KDM3A, basic helix-loop-helix factors MYC, HIF1A, and SREBF1, as well as differentiation factors AP1, MYOD1, SP1, MEIS1, ZEB1 and ELK1. In such a cooperation network, one component opens the chromatin, another one recognizes gene-specific DNA motifs, others scaffold between histones, cofactors, and the transcriptional complex. In cancer, due to the ability to team up with transcription factors, epigenetic factors concert mitogenic and metabolic gene networks, claiming the role of a cancer master regulators or epioncogenes.

Specific histone modification patterns are commonly associated with open or closed chromatin states, and are linked to distinct biological outcomes by transcriptional activation or repression. Disruption of patterns of histone modifications is associated with loss of proliferative control and cancer. There is tremendous therapeutic potential in understanding and targeting histone modification pathways. Thus, investigating cooperation of chromatin remodelers and the transcriptional machinery is not only important for elucidating fundamental mechanisms of chromatin regulation, but also necessary for the design of targeted therapeutics.

## Introduction

Beyond genomic alterations, aberrant epigenomes contribute to many cancers, as demonstrated by widespread changes to DNA methylation patterns, redistribution of histone marks and disruption of chromatin structure ^1^. Altered epigenomes and transcriptomes are closely intertwined and share non-genomic mechanisms of dysregulation in cancer, and are therefore not just a passive by-product of cancer ^2^. Epigenomic modifiers have the ability to affect the behavior of an entire network of cancer genes and can take on oncogenic roles themselves ^3^. Furthermore, epigenetic factors cooperate and team up with transcription factors to control specific gene target networks ^4,5^. In such works and in the following text, a *cis*-regulatory, synergistic molecular event between epigenetic and transcription factors is referred to as transcriptional cooperation (Figure 1).

**Figure 1:**
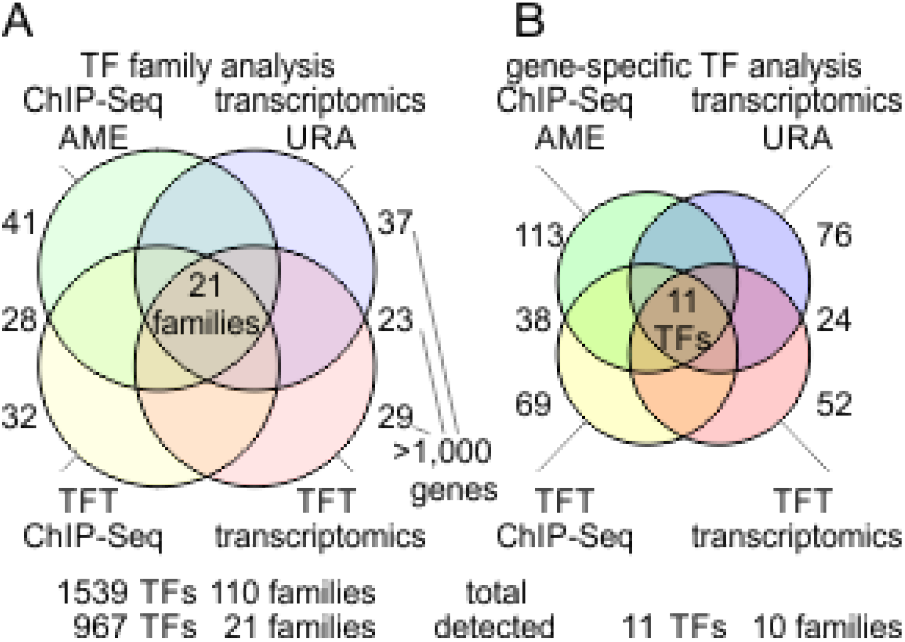
Transcription factor enrichment associated with activity of epigenetic modifier assessed by different omics platforms and analysis techniques at the family and gene level.

The combination of both, transcriptomic and epigenomic profiling offers insight into different levels of gene regulation, transcription factor binding motifs, DNA and chromatin modifications, and how each component is coupled to a functional output. Chromatin remodelers and transcription factors are in close communication via recognition of post-translational histone modifications ^6^. Thereby, they have the ability to harmonize and synchronize a dynamic exchange of chromatin between open, transcriptionally active conformations and compacted, silenced ones ^3^. Coordinated experiments interrogating transcriptional responses and chromatin binding via chromatin immunoprecipitation with next generation sequencing (ChIP-Seq) are well-equipped to capture different epigenomic and transcriptomic levels governing the circuitry of a regulatory network ^5^.

Regulatory networks in biology are intrinsically hierarchical and governed by interactions and chemical modifications ^7,8^. The regulome describes the interplay between genes and their products and defines the control network of cellular factors determining the functional outcome of a genomic element. The reconstruction of regulatory gene networks is stated as one of the main objectives of systems biology ^9,10^. However, an accurate description of the regulome is a difficult task due to the dynamic nature of epigenetic, transcriptional, and signaling networks. Systems biology has the ability to integrate genome-wide omics data recorded by ChIP-Seq, assay for transposase-accessible chromatin using sequencing (ATAC-Seq), whole genome bisulfite sequencing (WGBS-Seq), and RNA sequencing (RNA-Seq) technology to identify gene targets of a regulatory event ^11^. The integrated analysis of such data―on the one hand based on gene networks, on the other hand based on sequence features of high-resolution sequencing data—captures cooperation among regulators. Effective experimental design and data analysis of complementary epigenomic and transcriptomic platforms are required to decipher such epigenomic and transcriptional cooperation, which has profound impact in development and disease.

We took advantage of published, information-rich transcriptomic and epigenomic data to study regulatory networks surrounding histone lysine demethylation. The presence or absence of methylation on histone lysine residues correlates with altered gene expression and is an integral part of the epigenetics code ^12^. In particular histone 3 lysine 9 methylation (H3K9) is regarded as an epigenetic mark associated with suppressed gene expression ^13^. The H3K9 lysine demethylase 3A (KDM3A, also referred to as JMJD1A, Gene ID: 55818, HGNC ID: 20815) demethylates mono- or di-methylated histone marks, thereby influencing gene regulation within spermatogenesis, metabolism, stem cell activity and tumor progression ^14–16^. Genome-wide ChIP-Seq data of KDM3A identified specific gene targets and transcriptional networks in androgen response, hypoxia, glycolysis, and lipid metabolism, emphasizing the importance of cooperation with transcription factors ^5^. However, among epigenetic profiling experiments, a common observation is that enrichment studies provide significance for multiple transcription factors and not just one single, prioritized hit. This underscores the concept of transcriptional cooperation among epigenetic players but also emphasizes the need to design a reliable workflow that includes cross-validation with complementary, multi-omics platforms and analysis techniques.

## Results

### Deciphering the regulatory landscape of an epigenomic and transcriptomic network

The regulatory landscape of an epigenomic player includes histone modifications, non-enzymatic chromatin interactions, cooperation with transcription factors, transcriptional modulation of gene target networks, and eventually stimulation of specific effector pathways. With the help of hierarchical experimental design, the complementary power of epigenomic and transcriptomic data can be leveraged, and thus used to address distinct levels of the regulome ^4^. However, different genome-scale data platforms and analysis techniques result in the detection of significant, yet only occasionally overlapping, insight into regulatory networks. To address this, the intersection of multi-omics data levels is useful to augment and validate epigenetic regulation of transcriptional programs. At the same time, there are unique, platform-specific insights, which need to be analyzed accordingly.

### Prioritization of cooperating transcription factors by integration of complementary data

The goal of our network analysis is to detect epigenomic and transcriptomic cooperation. Specifically, it is of interest to identify and prioritize transcription factors that are closely associated with an epigenomic factor by integrating complementary data (Figure 1). Analysis of motif enrichment (AME) determines significant enrichment of transcription factor motifs among promotors or given sequences. Transcription factor target (TFT) analysis defines which transcription factor governs a set of target genes. Upstream regulator analysis (URA) integrates TFT networks with reconstructions of systems biology maps. Each platform provides a measure of significance for each detected transcription factor feature and corrects for multiple hypothesis testing using unbiased genome-wide data ^17^. Importantly, computations of significance of enrichment may be performed at the level of either the gene or the sequence. Furthermore, different members of transcription factor families have the ability to recognize the same sequence motif. Therefore, dedicated searches may account for such ambiguity and overcome potential gene-specific database biases of individual transcription factors (Figure 2). In a subsequent step, results of complementary platforms are intersected and compared. Taken together, combinations of high-throughput sequencing data deliver coordinates of epigenomic modification, enrichment of transcription factor motifs, transcriptional output, and networks of transcription factor targets (Figure 1 and 2).

**Figure 2:**
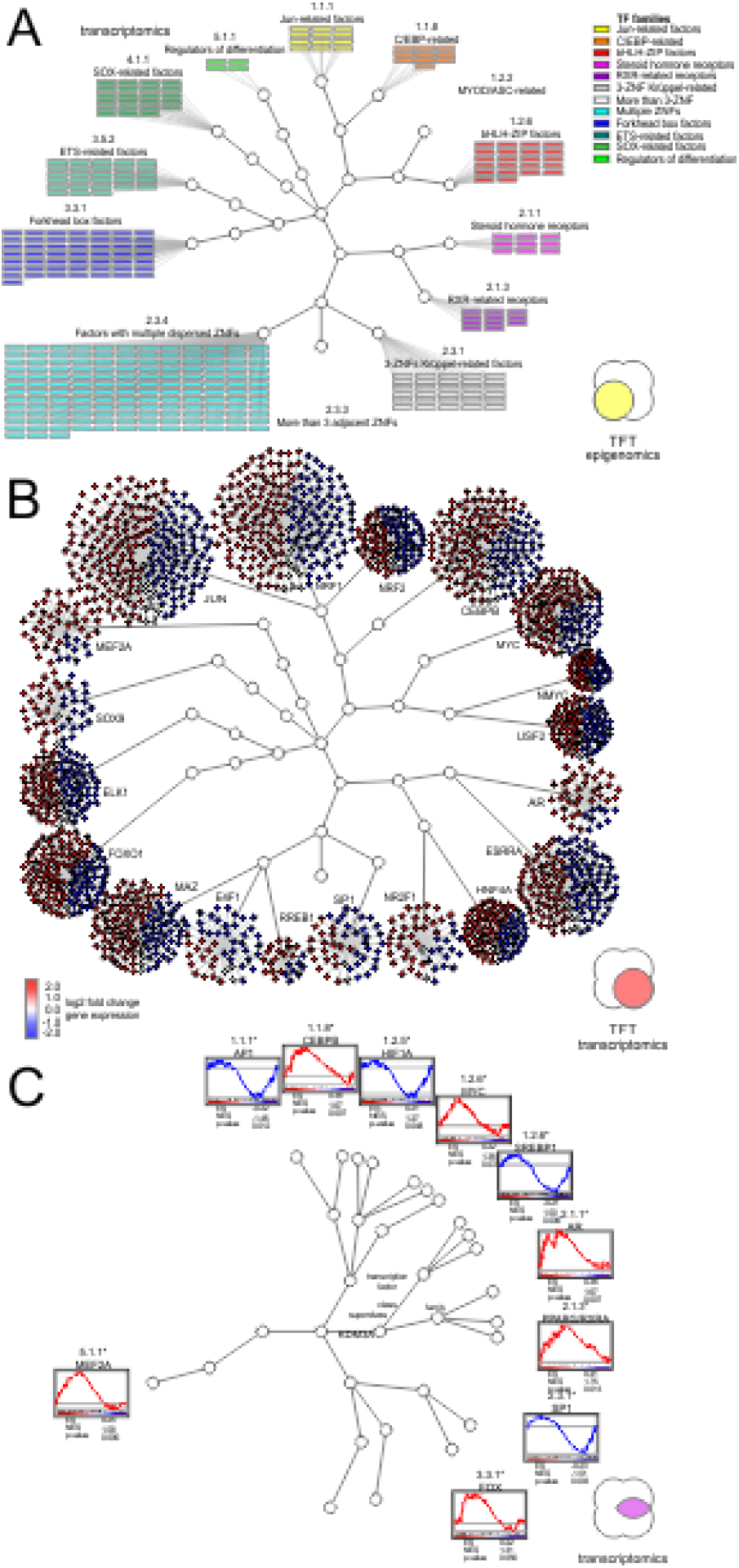
Visualization of epigenomic and transcriptional cooperation illustrates redundancy and complexity of a target network.

### Epigenomic switch of histone demethylation makes chromatin accessible and activates gene expression

Our approach is showcased by elucidating the epigenomic switch of KDM3A emphasizing its role as a master regulator. In order to better understand the impact of KDM3A on transcriptional networks, coordinated ChIP-Seq and transcriptomic data of KDM3A binding and demethylation activity in combination with knockdown of KDM3A was utilized ^5^. Such a combined array of matching epigenomics and transcriptomics experiments has the ability to focus on the cooperative forces of epigenetic regulation as well as its transcriptional consequences. A ChIP-Seq experiment offers direct insight into chromatin binding events and chemical modifications of histones. By overlaying genomic binding events with tracks of epigenomic marks, such as histone acetylation or methylation, associated with open or closed states of chromatin, a functional epigenomic landscape arises. Such ChIP-Seq profiles in combination with transcriptomics and functional genomics allow interrogation of the genome-wide impact of knockdown of a specific epigenomic regulator. Via genome-wide annotation and integration of sequencing reads, it becomes apparent that corresponding profiles of binding and histone modifications are reversed upon loss of function, mirroring the enzymatic function of the epigenetic modifier. Cooperative epigenomic and transcription factor binding coincides with promoter sites on meta gene coordinates enriched for histone lysine demethylation―overall indicators of transcriptionally activating epigenetic remodeling.

### Regulation of transcriptional networks by H3K9 chromatin demethylation

Coordinated ChIP-Seq and transcriptomic data classified genome-wide interactions of the chromatin demethylase KDM3A using antibodies specific for KDM3A, and its histone marks H3K9me1 and H3K9me2, conjointly with shRNA knockdown of KDM3A in the CWR22Rv1 cell line. Transcriptomic impact of 4326 differentially expressed genes upon KDM3A knockdown showed 2460 genes as positively regulated by KDM3A activity (down in the prostate cancer line CWR22Rv1 with shRNA knockdown of KDM3A), and 1866 genes as negatively regulated by KDM3A activity. Using this data we defined the set of 56.9% differentially expressed genes as positively regulated by KDM3A activity (down in the prostate cancer line CWR22Rv1 with shRNA knockdown of KDM3A), and 43.1% of differentially expressed genes as negatively regulated by KDM3A activity. KDM3A binding locations were defined by a loss of ChIP-Seq signal following knockdown of KDM3A. Concurrently, H3K9me1/2 histone marks following KDM3A knockdown are recognized as target regions of KDM3A histone lysine demethylation mediated by KDM3A. Changes in these ChIP-Seq marks upon KDM3A knockdown were contrasted against reference genomic DNA input or control non-coding shRNA samples. Quantification of the activity-based ChIP-Seq array matched with knockdown of KDM3A resulted in 37525 peaks associated with KDM3A binding, 45246 and 32665 H3K9 mono- and di-demethylation (H3K9me1/2-KDM) events, respectively. Overall, the peak counts of both histone marks showed a gain of signal upon knockdown of KDM3A reflecting the demethylase activity. By integrating continuous ChIP-Seq signals, an average profile of a meta-gene can be generated and functional coordinates analyzed for regulatory control. In such a meta-gene profile, promotor regions are located within 1000bp upstream of the gene-coding body, with the transcription start site (TSS) as the start of the gene-coding body at the zero position, and intergenic regions as the remaining regions outside of the gene body. KDM3A localized to the response element-rich promoter regions and demethylated H3K9me1/2 histone marks in the proximity of the TSS. Taken together, the meta-gene analysis classified areas important for transcriptional regulation and defined genomic sequence coordinates relevant for cooperation with transcription factors.

### Accounting for motif similarity and structural homology of transcription factor families

The array of ChIP-Seq data was subjected to AME and TFT analysis, while the list of differentially expressed genes served as input for URA and TFT analysis (Figure 1A). Each transcription factor hit was reported with its HGNC identification number, hierarchical classification of human transcription factors (TFClass) family barcode, motif logo, and significance corrected for multiple hypothesis testing using an adjusted *p* value cut-off of 0.05 (Table 1, Supplementary Information 1). Each individual analysis yielded between 29 and 41 significantly enriched transcription factor families, corresponding to 1083 up to 1292 associated genes (Figure 1B). In comparison to the entire realm of 1539 existing transcription factors, such wide-ranging data tables provide little benefit, despite the impressive *p* values produced by analysis tools at first glance. For example hypoxia inducible factor 1 alpha subunit (HIF1A, Gene ID: 4609, HGNC ID: 4910, TFClass: 1.2.5) of the PAS domain factors (TFClass 1.2.5) is detected with an adjusted *p* value below 1.0E-100 by AME in the ChIP-Seq data. Other members of the same family like the aryl hydrocarbon receptor nuclear translocator (ARNT, Gene ID: 405, HGNC ID: 700, TFClass: 1.2.5) show similar significance, since the detection is based on the same sequence logo, highlighting the lack of ability to differentiate between structurally homologous transcription factors. Therefore, we intersected all four sets of AME ChIP-Seq, TFT ChIP-Seq, URA transcriptomics, and TFT transcriptomics, and narrowed down 21 transcription factor families supported by all datasets (Figure 1A). Despite a considerable improvement of 21 projected families among 110 existing transcription factor families, the final set maps back to 967 transcription factors. In part, such lack of specificity is due to the large family of more than 3 adjacent zinc finger factors (TFClass: 2.3.3), whose motif was detected by the analysis but contains 487 members, accounting for almost a third of all transcription factors (Figure 2A). Systems biology networks and enrichment studies provide insight into directionality of the response and draw attention to different sized effector networks (Figure 2B-C). Only few nodes of the transcription factor target network were hyperconnected and showed promoter association with multiple transcription factors in epigenomics and transcriptomics datasets (Figure 1 and 2). Such a high degree of network connectivity speaks to a synergistic effect, where selected master regulators cooperate and act in sync, resulting in robust transcriptional output.

**Table 1:**
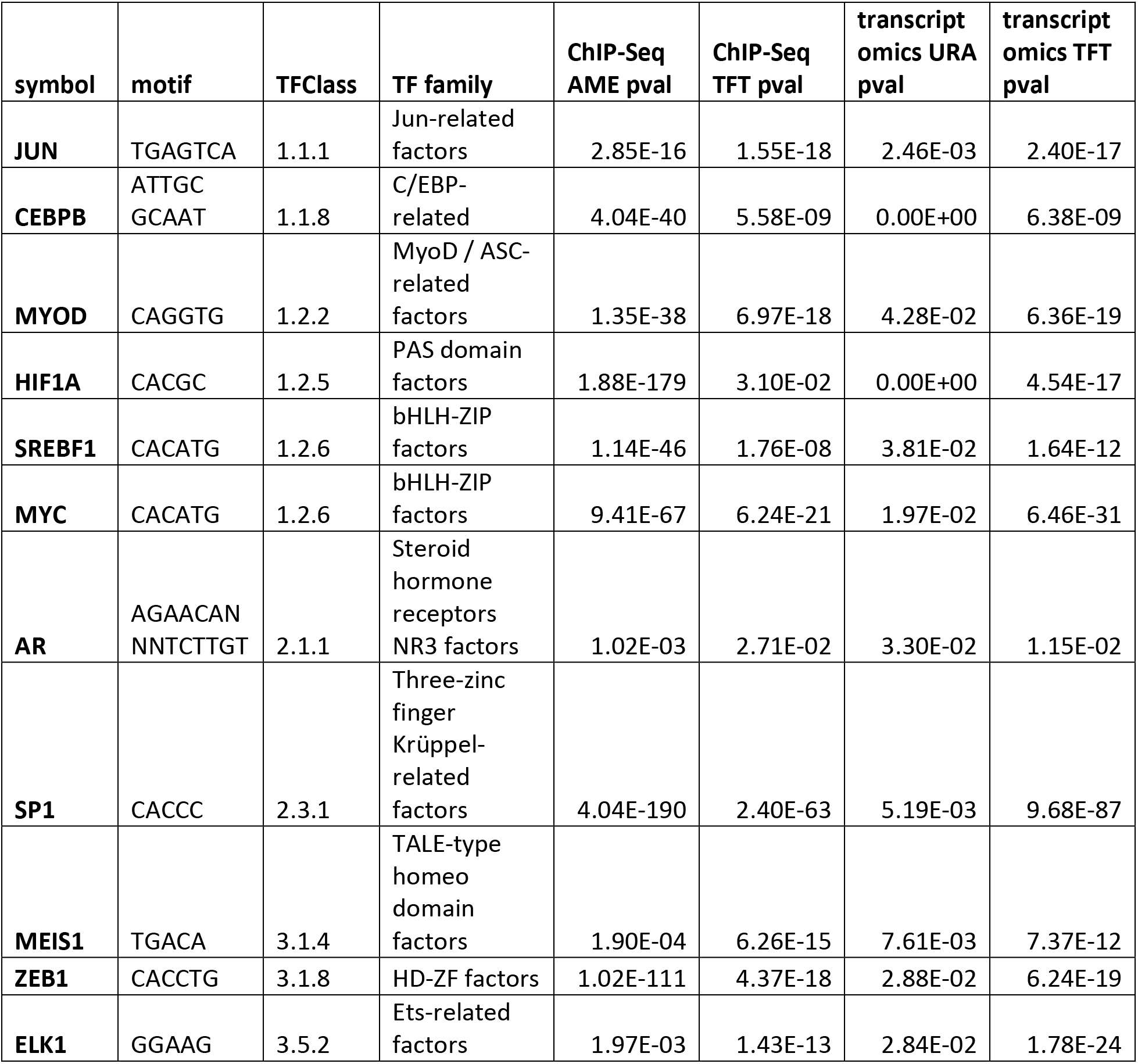
Detection of transcriptional cooperation by multi-omics integration of complementary data and analysis techniques. ChIP-Seq or transcriptomics data provide adjusted *p* values (pval) using analysis of motif enrichment (AME), transcription factor target analysis (TFT), or up-stream regulator analysis (URA).

### Multi-omics integration of complementary data yields well-refined target network

In order to improve the detected output, gene-specific systems biology networks were employed. In particular, URA and TFT analysis fueled by transcriptomic data provide useful insight. Databases of gene sets rely in part on experimental data of gene-specific knockdowns to characterize the impact of a transcription factor on a target network. Furthermore, a consistent directional response amplifies the significance of a detected hit. Therefore, such directional, gene-specific networks have the ability to overcome ambiguities. For example, members of the JUN-related factors (TFClass: 1.1.1) are detected but show different signs of regulation depending on the factor of interest, the response of its target genes, and the change of expression of the factor itself. After intersection of all four datasets, 11 transcription factors belonging to 10 transcription factor families were determined (Figure 2C). This set of master regulators is supported by complementary omics platforms and different analysis techniques representing a high-confidence cooperation network of the epigenomic master regulator KDM3A (Figure 1 and 2). The cooperation network includes cancer associated factors Jun proto-oncogene, AP-1 transcription factor subunit (JUN, Gene ID: 3725, HGNC ID: 6204, TFClass: 1.1.1), CCAAT/enhancer binding protein beta (CEBPB, Gene ID: 1051, HGNC ID: 1834, TFClass: 1.1.8), myogenic differentiation 1 (MYOD1, Gene ID: 3091, HGNC ID: 7611, TFClass: 1.2.2), HIF1A, sterol regulatory element binding transcription factor 1 (SREBF1, Gene ID: 379, HGNC ID: 11289, TFClass: 1.2.6), MYC proto-oncogene, bHLH transcription factor (MYC, Gene ID: 4609, HGNC ID: 7553, TFClass: 1.2.6), androgen receptor (AR, Gene ID: 367, HGNC ID: 644, TFClass: 2.1.1), Sp1 transcription factor (SP1, Gene ID: 6667, HGNC ID: 11205, TFClass: 2.3.1), Meis homeobox 1 (MEIS1, Gene ID: 4211, HGNC ID: 7000, TFClass: 3.1.4), zinc finger E-box binding homeobox 1 (ZEB1, Gene ID: 6935, HGNC ID: 11642, TFClass: 3.1.8), and ELK1, ETS transcription factor (ELK1, Gene ID: 2002, HGNC ID: 3321, TFClass: 3.5.2) representing less than 0.8 percent of all possible transcription factors (Figure 1B and 2A). The identified factors can further be surveyed by basal expression level or regulation in patient-derived tumor sample of TCGA underlining elevated expression in tumor progression. The analysis validates previously reported associations and contacts implicated in chromatin remodeling and discovered newly identified cooperative interactions (Table 2). For the specific role of KDM3A in cancer, epigenomic and transcriptional cooperation with transcription factors is key. KDM3A cooperates with mitogenic basic helix-loop-helix factors including MYC, HIF1A, and SREBF1, and derives a lipogenic program from association with nuclear receptors like AR. Ultimately, by overlaying the motif-specific and genomic data produced through matched experiments, epigenomic events can be correlated with the transcriptomic effect of histone remodelers and transcription factors.

**Table 2:** Epigenomic and transcriptional cooperation events in cancer. Original findings and reported cooperation events of epigenomic regulators with transcription factor are enumerated.

| Symbol | TFClass | Cooperation event | Ref. |
| --- | --- | --- | --- |
| JUN | 1.1.1 | KDM3A assists in recruiting JUN to AP1 binding sites in regulating expression of CD44, MMP7, and PDGFRB in liver adenocarcinoma tumor formation. | 45 |
|  |  | KDM4A promotes a positive feedback loop by facilitating the binding of the AP1 complex to the promoters JUN and FOSL1 in squamous cell carcinoma cells. | 46 |
| CEBPB | 1.1.8 | Novel event |  |
|  |  | KDM4B serves as a cofactor for CEBPB in preadipocytes and is recruited to the promoters of CEBPB regulated cell cycle genes. | 47 |
| MYOD | 1.2.2 | Novel event |  |
|  |  | KDM4B regulates the expression of MYOD and physically interacts with MYOD thereby controlling myogenic differentiation. | 48 |
|  |  | KDM4C decreases MYOD degradation and increase MYOD transcriptional activity to facilitate skeletal muscle differentiation. | 49 |
| HIF1A | 1.2.5 | KDM3A expression is stimulated by HIF1A binding to a response element in the promoter region of KDM3A. | 24 |
|  |  | KDM3A is regulated by HIF1A stimulating tumor formation in renal cell carcinoma. | 22 |
|  |  | KDM3A cooperates with HIF1A to induce glycolytic genes in urothelial bladder carcinoma. | 25 |
| SREBF1 | 1.2.6 | Novel event |  |
|  |  | KDM1A regulates SREBF1 binding to the FASN promotor stimulating lipogenesis. | 26 |
| MYC | 1.2.6 | KDM3A stimulates MYC expression and attenuates its ubiquitin-dependent degradation by binding to a E3 ubiquitin ligase. | 50 |
|  |  | KDM3A regulates transcription of MYC and PAX3 by directly binding to their promoters and regulates their H3K9me2 level in breast adenocarcinoma. | 51 |
|  |  | KDM4B binds the MYC/MAX motif and regulates expression of MYC signaling in neuroblastoma | 52 |
|  |  | N-MYC physically interacts and recruits KDM4B. Additionally KDM4B is able to regulate the expression of MYC signaling in neuroblastoma | 52 |
| AR | 2.1.1 | KDM3A regulates the transcriptional program of the AR, serves epigenomic master regulator by epigenomic and transcriptional cooperation of prostate adenocarcinoma. | 5 |
|  |  | KDM3A facilitates transcriptional activation by hormone-dependent recruitment of the AR to target genes in prostate adenocarcinoma. | 14 |
|  |  | KDM1A and KDM4D bind to the AR and localize to ARE half sites in the promoter region of VEGFA in placental development. | 21 |
|  |  | KDM4A binds the AR and supporting urothelial bladder carcinoma initiation and progression. | 53 |
|  |  | KDM4B enhances AR transcriptional activity by demethylation and inhibits ubiquitination of the AR. | 54 |
| SP1 | 2.3.1 | Novel event |  |
|  |  | KDM4A silences SP1 by chromatin demethylation in breast adenocarcinoma. | 55 |
| MEIS1 | 3.1.4 | Novel event |  |
| ZEB1 | 3.1.8 | Novel event |  |
| ELK1 | 3.5.2 | Novel event |  |

## Methods

### Experimental design

Coordinated epigenomic and transcriptomic profiles are well-equipped to capture different regulatory levels governing the circuitry of a cooperative network. The workflow introduced in our approach is open and applicable to different epigenome and transcriptome profiles including ChIP-Seq, ATAC-Seq, RNA-Seq, and microarray experiments. Epigenomic modifiers coordinate transcriptional activation by histone and DNA modifications thereby enabling chromatin accessibility and replacement of components of nucleosome-stalled polymerase complexes by tissue-specific transcription factors. The workflow is exemplified by reprocessing published data on KDM3A activity recorded with matching knockdown conditions in a human prostate carcinoma epithelial cellular model deposited in NCBI GEO entries GSE109748 and GSE70498 ^29,36,37^ (CRL-2505, American Type Culture Collection, Manassas, VA). The utilized cellular model is a variant derived from a xenograft and simulates castration-induced regression and relapse typical of human prostate carcinoma epithelial cells independent of dihydroxytestosterone stimulation. Furthermore, KDM3A is activated by somatic copy number amplification in lung, prostate, uterine, bladder, testicular germ cell, ovarian, cervical, breast, sarcoma, melanoma, and other cancers making its cooperation network an important target of broad interest in oncology.

### Data processing

Illumina HiSeq 2000 (Illumina, San Diego, CA) fastq files were aligned to the reference human genome 19 using the Bowtie software package ^38^. Peak-calling utilized a model-based analysis of ChIP-Seq (MACS) algorithm ^39,40^. Significant ChIP-Seq genomic locations relative to nearby gene bodies were annotated by ChIPSeek ^41^. ChIP-Seq peak regions were sorted and filtered by BEDtools ^42^. Average ChIP enrichment profiles over specific genomic features were calculated using the Cis-regulatory element annotation system tool ^43,44^. ChIPSeq binding profiles were visualized in the integrative genomics viewer (IGV) ^45^. Utilized conditions include ChIP-Seq profiles of antibodies specific for chromatin marks H3K9me1, H3K9me2, and the epigenomic modifier KDM3A in combination with small hairpin RNA (shRNA) knockdown of KDM3A matched with coordinated transcriptomic data of control and KDM3A knockdown cells using human transcriptome platform GPL10558 (HT-12 V4.0, Illumina, San Diego, CA) ^5^. The epigenomic and transcriptomics datasets contained 77911 features and 4356 differentially expressed transcripts upon KDM3A knockdown, respectively, with *p* values and *q* values below 0.05 adjusted for multiple hypothesis testing.

### Network analysis and transcription factor target enrichment

Human transcription factors were annotated according to their Human Genome Organization (HUGO) Gene Nomenclature Committee (HGNC) identification number using the using the multi-symbol checker tool. Discovered transcription factors were classified by shared DNA binding domains according to the hierarchical classification of human transcription factors (TFClass) database ^46^. Transcription factor binding and promoter sites were annotated utilizing transcription factor databases ^30,47^. For transcription factor binding site searches we built manual or utilized deposited position site-specific matrices or sequence logos of curated, non-redundant transcription factor databases. Statistically significant enrichment of these transcription factor motifs was determined using find individual motif occurrences (FIMO) and motif enrichment tools of the motif based sequence analysis toolkit (MEME) suite ^48^. Upstream regulators were determined by ingenuity pathway analysis (IPA, Qiagen, Redwood City, CA) based on differentially expressed genes with an adjusted *p* value below 0.05. Significant enrichment of target gene networks with transcription factor motif was calculated for target genes with annotated transcription factor motif in the 3’ promoter region of the transcription start site ^49^.

## Discussion

ChIP-Seq based approaches provide sequence resolution but detection of enriched transcription factor motifs is ambiguous and is most appropriately accomplished at the transcription factor family level to account for and include homologous factors. In contrast, transcriptomics studies provide directionality of regulation—transcriptional activation or repression upon epigenomic activity—an important aspect lacking in coordinate-based ATAC-Seq or ChIP-Seq experiments. Integration of different sequence, gene, or network-based approaches prioritizes high-fidelity cooperation partners in epigenomic regulation. Therefore, any combination of complementary data from sequence, gene, or network-based approaches is identified as desirable input for reliable regulatory systems biology analyses.

For the oncogenic nature and target specificity of an epigenomic master regulator, epigenomic and transcriptional cooperation with transcription factors is key. KDM3A is able to support initiation of transcription by its ability to specifically remove mono- and di-methylation marks from the H3K9 residue leading to chromatin decondensation ^14^. Transcriptionally silenced genes contain methylation marks on the H3K9 subunit^18^ and ChIP-Seq experiments monitoring H3K9 methylation marks showed global histone demethylation effects of KDM3A. Combined assessment of histone demethylation events and gene expression changes indicated major transcriptional activation, suggesting that distinct oncogenic regulators, in particular transcription factors, may synergize with the epigenetic patterns controlled by KDM3A. Furthermore, the epigenetic factor was shown to cooperate with the androgen receptor to control prostate tissue-specific gene target networks introducing the concept of the epioncogene and transcriptional cooperation ^19^.

While KDM3A is able to control chromatin accessibility, the mechanism by which it targets specific genes is of current interest and may influence understanding of epigenetic dysregulation in human disease. While several cancers exhibit deregulated KDM3A activity, in prostate adenocarcinoma it functions as a transcriptional coregulator with the androgen receptor ^5,14,20,21^. Such cooperative coactivation of the androgen receptor with KDM3A features a role for KDM3A as an active force in commencing oncogenesis in prostate epithelial cells. KDM3A is known to control the transcription and function of oncogenic transcription factors ^22,23^. However, an expanded study outlining the effects of perturbed KDM3A H3K9 demethylation upon human transcription factor response element recognition in cancer has so thus been missing.

The cooperation network includes previously validated interactions of MYC, HIF1A, and AR in cancer but also highlights close association of KDM3A with transcriptional networks of factors rather studied in development and tissue differentiation like JUN, CEBPB, MYOD, SREBF1, SP1, MEIS1, ZEB1, or ELK1 (Table 1). By taking advantage of motif-specific target networks, KDM3A has the ability to induce glycolytic genes in urothelial bladder carcinoma^24,25^. Epigenomic regulation of SREBF1 activity has been reported to stimulate lipogenesis, and SREBP1 regulates lipid accumulation and cell cycle progression in androgen independent prostate cancer cell lines ^26–28^. KDM3A regulates the transcriptional program of the AR, serves epigenomic master regulator by epigenomic and transcriptional cooperation of prostate adenocarcinoma ^5^. Despite some factors including the forkhead box (FOX) factors (TFClass: 3.3.1) family were frequently detected at the transcription factor family levels, lack of consistent overlap of epigenomic and transcriptomic data eventually excluded prominent cancer drivers like forkhead box A1 (FOXA1, Gene ID: 3169, HGNC ID: 5021, TFClass: 3.3.1), forkhead box M1 (FOXM1, Gene ID: 2305, HGNC ID: 3818, TFClass: 3.3.1), forkhead box O1 (FOXO1, Gene ID: 2308, HGNC ID: 3819, TFClass: 3.3.1), or forkhead box O3 (FOXO3, Gene ID: 2309, HGNC ID: 3821, TFClass: 3.3.1). Despite FOX factors are known to cooperate with nuclear hormone receptors (TFClass: 2.1.1), it is possible that the putative association with KDM3A steams from the fact that the closely cooperating AR frequently has FOX motifs nearby. Systematic, genome-wide surveys have elucidated that FOX motifs are adjacent androgen response elements (AREs) ^29,30^, thereby facilitating cooperation at the level of transcription factors and promotion of prostate cancer progression.

Hundreds of transcription factors are significantly associated with each individual data analysis platform or high-throughput sequencing technology. Big data challenges can be overcome by systems biology analysis and integration of multi-omics data. Motif similarity is visualized by transcription factor family trees classifying superclass, class, and family of transcription factors (from inward to outward, Figure 2) based on the characteristics of their DNA-binding domains. Single epigenomic or transcriptomic datasets examined by different analysis tools result in improved resolution but leave ambiguities. The footprints of cooperating transcription factors are found in cognate sequence motifs specific to their DNA binding domains. Such sequence motifs are more pronounced in events of cooperating epigenomic activity. Detected motif enrichment highlights the modularity, versatility, and efficacy of epigenomic cooperation, providing target specificity at genome-wide reach. The number of detected events in genome-wide epigenomic binding studies provides statistical power for sequence motif discovery and gene target enrichment. As a consequence, high-resolution epigenomic studies often arrive at multiple plausible solutions, though each suggested interaction or association may carry statistical significance. By intersecting complementary data platforms and analysis techniques, high-fidelity gene target networks involved in epigenomic and transcriptomic cooperation can be identified.

Within the regulome, epigenetic master regulators position themselves at the top of cellular hierarchies and control distinct phenotypic programs via reversible chemical modifications of chromatin, histone or nucleotide marks, without altering the core DNA sequence. Epigenetic oncogenes or tumor suppressors can arise when epigenetic master regulators are somatically activated or lost, and contribute to cancer initiation and progression ^31^. In cancer, such epigenetic master regulators are found at the top of regulatory hierarchies, particularly in pathways related to cellular proliferation, survival, fate, and differentiation. For the manifestation of a genomic or non-genomic aberration of an epigenetic master regulator, it is a necessity that its own activity is affected by somatic mutation, copy number alteration, expression levels, protein cofactors, or methylation status. Epigenetic master regulators often accomplish target specificity of their phenotypic program by cooperation with members of the transcriptional machinery and therefore may depend on tissue-specific expression of such auxiliary factors. In cancer, an epigenetic master regulator populates an extreme state and is either permanently switched on or off. An epigenetic master regulator will become a cancer driver, if it is not functionally neutral but rather contributes to tumorigenesis or disease progression due to its hyperactive or deactivated state. Genomic profiling of cancer patients has the ability to identify coincidence or mutual exclusivity of somatic alterations of epigenomic and transcription factors. Extreme states of epigenetic master regulators by somatic loss or gain of function in cancer may emphasize preexisting cooperative interactions with transcription factors, which may be subtle and difficult to detect under normal circumstances. A defined challenge in the field of epigenetic master regulators is to identify cancer-specific vulnerabilities in gene targets and biological pathways that are frequently and consistently perturbed under the control of an epigenetic driver.

## Conclusion

Epigenetic regulators such as KDM1A, KDM3A, KDM5A, KDM6A, KDM7A, EZH2, DOT1L, and others have been shown to be critical in oncogenesis and cancer resistance ^3,32^. The identification of transcriptional cooperation and regulatory hierarchies highlights the importance of epigenetic regulators in mitogenic control and their potential as therapeutic targets ^33^. The discovery of the specific role of KDM3A in the interplay between a tissue specific steroid receptor transcription factor and metabolic signaling provides a foundation for rational design of combination approaches where metabolic, epigenetic, and hormone-deprivation therapies may synergize. Our integrated multi-platform analysis reveals a complex molecular landscape of epigenomic and transcriptomic cooperation in cancer, providing avenues for precision medicine ^34^. A close teamwork of the transcriptional and epigenomic machinery was discovered, in which one component opens the chromatin, another recognizes gene-specific DNA motifs, and others scaffold between histones, cofactors, and the transcriptional complex. This highlights a close connection between the epigenomic and transcriptomic machinery, albeit much of the underlying principles remain to be discovered. In conclusion, transcriptomics in combination with epigenomic profiling and measurement of chromatin accessibility enable global detection of epigenetic modifications and characterization of transcriptional and epigenetic footprints. Chromatin remodelers and transcription factors are in close communication via recognition of post-translational histone modifications and coordinate the dynamic exchange of chromatin between open, transcriptionally active conformations and compacted, silenced ones. In cancer, due to the ability to team up with transcription factors, epigenetic factors concert mitogenic and metabolic gene networks, claiming the role of a cancer master regulators or epioncogenes. Exploration into the cooperative roles of epigenetic histone modifiers and transcription factor families in gene regulatory networks contributes to our understanding of how a seemingly promiscuous epigenomic program is converted into a specific transcriptional response assisting in oncogenesis.

Hierarchical trees of human transcription factors correspond to transcription factor superclass, class, and family from inward out. Transcription factor motifs often get recognized by multiple members of the same transcription factor family due to structural homology of DNA binding domains. The transcription factor target analysis (TFT) is carried out on sequence-specific epigenomics data.

## Declarations

### Ethics approval and consent to participate

All experimental protocols were approved by the Institutional Review Boards at the University of California Merced. The study was carried out as part of IRB UCM13-0025 of the University of California Merced and as part of dbGap ID 5094 for study accession phs000178.v9.p8 on somatic mutations in cancer and conducted in accordance with the Helsinki Declaration of 1975.

### Availability of supporting data

Data is deposited in NCBI GEO entries GSE109748 and GSE70498.

### Preprint availability

A preprint version of this manuscript is made available to the scientific community on the preprint server BIORXIV/2018/309484 https://doi.org/10.1101/309484.

### Competing interest

The authors declare that there is no competing interest as part of the submission process of this manuscript.

### Author contribution statement

S.W. and F.V.F. conducted the data analysis. F.V.F. designed the network study, directed the systems biology analysis, performed data interpretation, prepared the illustrations, and wrote the text. S.W. and F.V.F. reviewed the final manuscript.

## Acknowledgements and funding

Science is breathing the creativity of life. This work on cancer systems biology is generously supported by the National Institutes of Health and the National Science Foundation. F.V.F. is grateful for the support of grant CA154887 from the National Institutes of Health, National Cancer Institute and grant CRN-17-427258 by the University of California, Office of the President, Cancer Research Coordinating Committee. Dedicated to Emilia Mili Filipp.

## References

1 Dawson, M. A. The cancer epigenome: Concepts, challenges, and therapeutic opportunities. Science 355, 1147-1152, doi:10.1126/science.aam7304 (2017).

2 Leung, J. K. & Sadar, M. D. Non-Genomic Actions of the Androgen Receptor in Prostate Cancer. Front Endocrinol (Lausanne) 8, 2, doi:10.3389/fendo.2017.00002 (2017).

3 Filipp, F. V. Crosstalk between epigenetics and metabolism-Yin and Yang of histone demethylases and methyltransferases in cancer. Brief Funct Genomics 16, 320-325, doi:10.1093/bfgp/elx001 (2017).

4 Filipp, F. V. Epioncogenes in cancer— Identification of epigenomic and transcriptomic cooperation-networks by multi-omics integration of ChIP-Seq and RNA-Seq data. Systems Biology. Methods Mol Biol 1800, 101-121 (2019).

5 Wilson, S., Fan, L., Sahgal, N., Qi, J. & Filipp, F. V. The histone demethylase KDM3A regulates the transcriptional program of the androgen receptor in prostate cancer cells. Oncotarget 8, 30328-30343, doi:10.18632/oncotarget.15681 (2017).

6 Ahsendorf, T., Muller, F. J., Topkar, V., Gunawardena, J. & Eils, R. Transcription factors, coregulators, and epigenetic marks are linearly correlated and highly redundant. PLoS One 12, e0186324, doi:10.1371/journal.pone.0186324 (2017).

7 Cheng, C. et al. An approach for determining and measuring network hierarchy applied to comparing the phosphorylome and the regulome. Genome Biol 16, 63, doi:10.1186/s13059-015-0624-2 (2015).

8 Ay, A., Gong, D. & Kahveci, T. Hierarchical decomposition of dynamically evolving regulatory networks. BMC Bioinformatics 16, 161, doi:10.1186/s12859-015-0529-9 (2015).

9 Filipp, F. V. Cancer metabolism meets systems biology: Pyruvate kinase isoform PKM2 is a metabolic master regulator. J Carcinog 12, 14, doi:10.4103/1477-3163.115423 (2013).

10 Filipp, F. V. A Gateway between Omics Data and Systems Biology. J Metabolomics Syst Biol 1, 1 (2013).

11 Gonda, T. J. & Ramsay, R. G. Directly targeting transcriptional dysregulation in cancer. Nat Rev Cancer 15, 686-694, doi:10.1038/nrc4018 (2015).

12 Zentner, G. E. & Henikoff, S. Regulation of nucleosome dynamics by histone modifications. Nat Struct Mol Biol 20, 259-266, doi:10.1038/nsmb.2470 (2013).

13 Kooistra, S. M. & Helin, K. Molecular mechanisms and potential functions of histone demethylases. Nat Rev Mol Cell Biol 13, 297-311, doi:10.1038/nrm3327 (2012).

14 Yamane, K. et al. JHDM2A, a JmjC-containing H3K9 demethylase, facilitates transcription activation by androgen receptor. Cell 125, 483-495, doi: 10.1016/j.cell.2006.03.027 (2006).

15 Okada, Y., Scott, G., Ray, M. K., Mishina, Y. & Zhang, Y. Histone demethylase JHDM2A is critical for Tnp1 and Prm1 transcription and spermatogenesis. Nature 450, 119-123, doi:nature06236 (2007).

16 Kuroki, S. et al. Epigenetic regulation of mouse sex determination by the histone demethylase Jmjd1a. Science 341, 1106-1109, doi:10.1126/science.1239864 (2013).

17 Storey, J. D. & Tibshirani, R. Statistical significance for genomewide studies. Proc Natl Acad Sci U S A 100, 9440-9445, doi:10.1073/pnas.1530509100 (2003).

18 Barth, T. K. & Imhof, A. Fast signals and slow marks: the dynamics of histone modifications. Trends Biochem Sci 35, 618-626, doi:10.1016/j.tibs.2010.05.006 (2010).

19 Qi, J. & Filipp, F. V. An epigenetic master regulator teams up to become an epioncogene. Oncotarget 8, 29538-29539, doi:10.18632/oncotarget.16484 (2017).

20 Yamada, D. et al. Role of the hypoxia-related gene, JMJD1A, in hepatocellular carcinoma: clinical impact on recurrence after hepatic resection. Ann Surg Oncol 19 Suppl 3, S355-364, doi:10.1245/s10434-011-1797-x (2012).

21 Cleys, E. R. et al. Androgen receptor and histone lysine demethylases in ovine placenta. PLoS One 10, e0117472, doi:10.1371/journal.pone.0117472 (2015).

22 Krieg, A. J. et al. Regulation of the histone demethylase JMJD1A by hypoxia-inducible factor 1 alpha enhances hypoxic gene expression and tumor growth. Mol Cell Biol 30, 344-353, doi:10.1128/MCB.00444-9 (2010).

23 Ohguchi, H. et al. The KDM3A-KLF2-IRF4 axis maintains myeloma cell survival. Nat Commun 7, 10258, doi:10.1038/ncomms10258 (2016).

24 Wellmann, S. et al. Hypoxia upregulates the histone demethylase JMJD1A via HIF-1. Biochem Biophys Res Commun 372, 892-897, doi:10.1016/j.bbrc.2008.05.150 (2008).

25 Wan, W. et al. Histone demethylase JMJD1A promotes urinary bladder cancer progression by enhancing glycolysis through coactivation of hypoxia inducible factor 1alpha. Oncogene 36, 3868-3877, doi:10.1038/onc.2017.13 (2017).

26 Abdulla, A. et al. Regulation of lipogenic gene expression by lysine-specific histone demethylase-1 (LSD1). J Biol Chem 289, 29937-29947, doi:10.1074/jbc.M114.573659 (2014).

27 Nambiar, D. K., Deep, G., Singh, R. P., Agarwal, C. & Agarwal, R. Silibinin inhibits aberrant lipid metabolism, proliferation and emergence of androgen-independence in prostate cancer cells via primarily targeting the sterol response element binding protein 1. Oncotarget 5, 10017-10033, doi:2488 (2014).

28 Huang, W. C., Li, X., Liu, J., Lin, J. & Chung, L. W. Activation of androgen receptor, lipogenesis, and oxidative stress converged by SREBP-1 is responsible for regulating growth and progression of prostate cancer cells. Mol Cancer Res 10, 133-142, doi:10.1158/1541-7786.MCR-11-0206 (2012).

29 Wilson, S., Qi, J. & Filipp, F. V. Refinement of the androgen response element based on ChIP-Seq in androgen-insensitive and androgen-responsive prostate cancer cell lines. Sci Rep 6, 32611, doi:10.1038/srep32611 (2016).

30 Kulakovskiy, I. V. et al. HOCOMOCO: towards a complete collection of transcription factor binding models for human and mouse via large-scale ChIP-Seq analysis. Nucleic Acids Res 46, D252-D259, doi:10.1093/nar/gkx1106 (2018).

31 Tiffen, J., Wilson, S., Gallagher, S. J., Hersey, P. & Filipp, F. V. Somatic Copy Number Amplification and Hyperactivating Somatic Mutations of EZH2 Correlate With DNA Methylation and Drive Epigenetic Silencing of Genes Involved in Tumor Suppression and Immune Responses in Melanoma. Neoplasia 18, 121-132, doi:10.1016/j.neo.2016.01.003 (2016).

32 Filipp, F. V. How cancer can become therapy-resistant—epigenetics might play a role. Sci Am 318, 1, doi:https://www.scientificamerican.com/article/how-cancer-can-become-therapy-resistant/ (2018).

33 Zecena, H. et al. Systems biology analysis of mitogen activated protein kinase inhibitor resistance in malignant melanoma. BMC Syst Biol 12, 33, doi:10.1186/s12918-018-0554-1 (2018).

34 Filipp, F. V. Precision medicine driven by cancer systems biology. Cancer Metastasis Rev 36, 91-108, doi:10.1007/s10555-017-9662-4 (2017).

35 Filipp, F. V., Rajewsky, N., Barciszewski, J. Systems biology RNA technologies―transcriptional and epigenomic cooperation. Systems Biology 1613, 161-191 (2018).

36 Pretlow, T. G. et al. Xenografts of primary human prostatic carcinoma. J Natl Cancer Inst 85, 394-398 (1993).

37 Sramkoski, R. M. et al. A new human prostate carcinoma cell line, 22Rv1. In Vitro Cell Dev Biol Anim 35, 403-409, doi:10.1007/s11626-999-0115-4 (1999).

38 Langmead, B., Trapnell, C., Pop, M. & Salzberg, S. L. Ultrafast and memory-efficient alignment of short DNA sequences to the human genome. Genome Biol 10, R25, doi:10.1186/gb-2009-10-3-r25 (2009).

39 Zhang, Y. et al. Model-based analysis of ChIP-Seq (MACS). Genome Biol 9, R137, doi:10.1186/gb-2008-9-9-r137 (2008).

40 Feng, J., Liu, T. & Zhang, Y. Using MACS to identify peaks from ChIP-Seq data. Curr Protoc Bioinformatics Chapter 2, Unit 2 14, doi:10.1002/0471250953.bi0214s34 (2011).

41 Chen, T. W. et al. ChIPseek, a web-based analysis tool for ChIP data. BMC Genomics 15, 539, doi:10.1186/1471-2164-15-539 (2014).

42 Quinlan, A. R. & Hall, I. M. BEDTools: a flexible suite of utilities for comparing genomic features. Bioinformatics 26, 841-842, doi:10.1093/bioinformatics/btq033 (2010).

43 Shin, H., Liu, T., Manrai, A. K. & Liu, X. S. CEAS: cis-regulatory element annotation system. Bioinformatics 25, 2605-2606, doi:10.1093/bioinformatics/btp479 (2009).

44 Ji, X., Li, W., Song, J., Wei, L. & Liu, X. S. CEAS: cis-regulatory element annotation system. Nucleic Acids Res 34, W551-554, doi:34/suppl_2/W551 (2006).

45 Thorvaldsdottir, H., Robinson, J. T. & Mesirov, J. P. Integrative Genomics Viewer (IGV): high-performance genomics data visualization and exploration. Brief Bioinform 14, 178-192, doi:10.1093/bib/bbs017 (2013).

46 Wingender, E., Schoeps, T., Haubrock, M., Krull, M. & Donitz, J. TFClass: expanding the classification of human transcription factors to their mammalian orthologs. Nucleic Acids Res 46, D343-D347, doi:10.1093/nar/gkx987 (2018).

47 Khan, A. et al. JASPAR 2018: update of the open-access database of transcription factor binding profiles and its web framework. Nucleic Acids Res 46, D260-D266, doi:10.1093/nar/gkx1126 (2018).

48 Bailey, T. L., Johnson, J., Grant, C. E. & Noble, W. S. The MEME Suite. Nucleic Acids Res 43, W39-49, doi:10.1093/nar/gkv416 (2015).

49 Subramanian, A. et al. Gene set enrichment analysis: a knowledge-based approach for interpreting genome-wide expression profiles. Proc Natl Acad Sci U S A 102, 15545-15550, doi:10.1073/pnas.0506580102 (2005).

50 Nakatsuka, T. et al. Impact of histone demethylase KDM3A-dependent AP-1 transactivity on hepatotumorigenesis induced by PI3K activation. Oncogene 36, 6262-6271, doi:10.1038/onc.2017.222 (2017).

51 Ding, X. et al. Epigenetic activation of AP1 promotes squamous cell carcinoma metastasis. Sci Signal 6, ra28 21-13, S20-15, doi:10.1126/scisignal.2003884 (2013).

52 Guo, L. et al. Histone demethylase Kdm4b functions as a co-factor of C/EBPbeta to promote mitotic clonal expansion during differentiation of 3T3-L1 preadipocytes. Cell Death Differ 19, 1917-1927, doi:10.1038/cdd.2012.75 (2012).

53 Choi, J. H., Song, Y. J. & Lee, H. The histone demethylase KDM4B interacts with MyoD to regulate myogenic differentiation in C2C12 myoblast cells. Biochem Biophys Res Commun 456, 872-878, doi:10.1016/j.bbrc.2014.12.061 (2015).

54 Jung, E. S. et al. Jmjd2C increases MyoD transcriptional activity through inhibiting G9a-dependent MyoD degradation. Biochim Biophys Acta 1849, 1081-1094, doi:10.1016/j.bbagrm.2015.07.001 (2015).

55 Fan, L. et al. Regulation of c-Myc expression by the histone demethylase JMJD1A is essential for prostate cancer cell growth and survival. Oncogene 35, 2441-2452, doi:10.1038/onc.2015.309 (2016).

56 Zhao, Q. Y. et al. Global histone modification profiling reveals the epigenomic dynamics during malignant transformation in a four-stage breast cancer model. Clin Epigenetics 8, 34, doi:10.1186/s13148-016-0201-x (2016).

57 Yang, J. et al. The role of histone demethylase KDM4B in Myc signaling in neuroblastoma. J Natl Cancer Inst 107, djv080, doi:10.1093/jnci/djv080 (2015).

58 Kauffman, E. C. et al. Role of androgen receptor and associated lysine-demethylase coregulators, LSD1 and JMJD2A, in localized and advanced human bladder cancer. Mol Carcinog 50, 931-944, doi:10.1002/mc.20758 (2011).

59 Coffey, K. et al. The lysine demethylase, KDM4B, is a key molecule in androgen receptor signalling and turnover. Nucleic Acids Res 41, 4433-4446, doi:10.1093/nar/gkt106 (2013).

60 Li, L. et al. JMJD2A-dependent silencing of Sp1 in advanced breast cancer promotes metastasis by downregulation of DIRAS3. Breast Cancer Res Treat 147, 487-500, doi:10.1007/s10549-014-3083-7 (2014).

